# Benchmarking of Quantum SVM and Classical ML Algorithms for Prediction of Therapeutic Proteins

**DOI:** 10.1101/2025.04.30.651419

**Authors:** Purva Tijare, Naman Kumar Mehta, Gajendra P. S. Raghava

**Affiliations:** Department of Computational Biology, Indraprastha Institute of Information Technology, Okhla Phase 3, New Delhi-110020, India

**Keywords:** Quantum machine learning, Quantum algorithms, Quantum support vector machine, Classical machine learning, Predictive Modelling, Therapeutic proteins, Protein prediction, Peptide prediction, Classification datasets, Benchmarking, QSVM vs CML, Bioinformatics

## Abstract

Over the past decade, quantum machine learning, particularly quantum support vector machines (QSVMs), has emerged as an optimistic alternative to classical machine learning (CML) techniques. This study rigorously benchmarks the performance of QSVM and CML-based models across four diverse datasets relevant to therapeutic proteins and peptides. Specifically, we evaluated these approaches for the prediction of B-cell epitopes (CLBtope), exosomal proteins (ExoPropred), hemolytic peptides (HemoPI), and toxic peptides (Toxinpred3). The maximum area under the receiver operating characteristic curve (AUC) for the CLBtope dataset achieved was 0.68 for QSVM and 0.82 for CML models. For the ExoPropred dataset, the maximum AUCs were 0.66 (QSVM) and 0.72 (CML). In contrast, both QSVM and CML models demonstrated high performance on the HemoPI dataset, yielding maximum AUCs of 0.95 and 0.98, respectively. Similarly, for the Toxinpred3 dataset, the maximum AUCs were 0.84 (QSVM) and 0.94 (CML). All models were evaluated using independent validation datasets not used during training. These results suggest that although CML currently demonstrates superior predictive capability for these tasks, the similar progression in performance indicates potential for future advancements in QSVM.

**Highlights:**

- Comparative study of QSVM and CML models on four bioinformatics datasets
- QSVM performance tries to approach CML in tasks involving hemolytic and toxic peptide prediction
- Independent validation confirms robustness of performance metrics
- Results highlight the potential of QSVMs as real-world quantum hardware continues to matures

## 1. Introduction

Quantum technology is evolving rapidly, opening up exciting opportunities for innovation and discovery.[1] Quantum machine learning (QML), at the crossroads of quantum computing and machine learning, holds the potential to transform industries and research areas.[2] By harnessing the power of quantum computing, QML introduces a new way of building machine learning models aiming to enhance their performance.[3][4] This is made possible through quantum subroutines that can boost the efficiency of traditional ML algorithms, offering a promising path toward more powerful and scalable solutions.[5][6]. Traditional machine learning has transformed various fields by powering predictive analytics, pattern recognition, and intelligent decision-making.[7][8] It builds on established computational frameworks, employing models such as decision trees, support vector machines (SVMs), and deep learning.[9] While these approaches are highly effective, they often become computationally demanding and uninterpretable, particularly when working with high-dimensional datasets.[10][11][12]

QML takes advantage of core quantum computing principles like entanglement, superposition, and quantum parallelism to create algorithms that are at least in theory, outperforming classical machine learning in case of speed and efficiency.[13] Emerging techniques like Variational Quantum Circuits (VQC), Quantum Support Vector Machines (QSVM), and Quantum Neural Networks (QNNs) show significant potential in theory. However, their real-world application is still limited by the challenges of today’s quantum hardware.[14][15] Integrating quantum computing into conventional workflows and solutions has the potential to boost computational efficiency and data processing power.[16] In this study, we take a hands-on approach, comparing quantum and classical machine learning models across four different datasets focused on binary classification. Specifically, we put Quantum Support Vector Machines (QSVM) up against classical methods like standard SVMs to see how they really stack up in real-world tasks.[17]

### 1.1 Overview of Machine Learning and Quantum Computing

Machine Learning is at the heart of modern artificial intelligence, driving advancements in healthcare, finance, and natural language processing.[18] Techniques such as Support Vector Machines, Decision Trees, Neural Networks, and Ensemble Methods in classical machine learning are optimized for efficient operation on traditional computing hardware.[19][20][21][22] Despite their proven effectiveness, these models often face difficulties in managing high-dimensional data, computational complexity, and scalability.[23] In contrast, quantum computing relies on a fundamentally different computational model, utilizing quantum mechanics principles such as superposition, entanglement, and quantum parallelism.[13] These distinct characteristics enable quantum computers to solve specific complex mathematical problems at an exponentially faster rate than classical systems. As quantum hardware advances, researchers are increasingly investigating methods to integrate quantum computing with machine learning, giving rise to the expanding domain of Quantum Machine Learning (QML).[24] Among QML techniques, Quantum Support Vector Machines have drawn significant interest as a quantum-powered extension of classical SVMs.[25] By utilizing quantum kernel methods and quantum feature spaces, QSVM aims to process data more efficiently, potentially offering an edge in high-dimensional classification problems.[26] However, the real impact of these advantages still depends on advancements in quantum hardware and further refinements in the underlying algorithms.[13]

### 1.2 Why QSVM? Potential Advantages Over Classical ML

Due to their effectiveness in handling classification tasks, Support Vector Machines (SVMs) have become a popular choice as they excel at finding optimal decision boundaries, even in complex, high-dimensional feature spaces.[21] However, classical SVMs can struggle with computational bottlenecks, especially when dealing with large datasets or computing non-linear kernel functions. As data complexity increases, these challenges can lead to significant processing overhead and inefficiencies, making it difficult to scale classical SVMs effectively.[27] Quantum Support Vector Machines (QSVM) aim to overcome these challenges by utilizing quantum kernel estimation, where a quantum computer computes the kernel function in an exponentially large Hilbert space.[28] Initially, QSVM has the potential to significantly accelerate performance by efficiently computing kernel functions, possibly resulting in exponential or polynomial gains in computational speed compared to classical SVMs. Furthermore, quantum feature maps enable the encoding of classical data into high-dimensional quantum states, improving pattern recognition capabilities.[26] In some cases, QSVM may also provide solutions to problems that are very computationally expensive for classical kernel evaluations, making it a promising alternative for tackling complex classification tasks.

Despite its potential advantages, QSVM faces significant practical challenges, including limitations in current quantum hardware, noisy qubits, and high error rates.[30] These obstacles raise questions about its practical effectiveness. Given these constraints, it remains unclear whether QSVM has the potential to surpass classical machine learning models in practical-world applications.[31] This study aims to bridge that gap by experimentally evaluating whether QSVM offers any measurable advantage over traditional ML techniques when applied to real-world datasets.

### 1.3 Importance of Benchmarking QSVM vs. Classical ML

As quantum computing rapidly progresses, Quantum Machine Learning (QML) has garnered considerable interest. However, there is still a lack of systematic empirical studies comparing QML models to their classical counterparts. While theoretical research suggests that QSVM could outperform classical SVMs in certain scenarios, its real-world effectiveness depends on several factors such as the nature of the dataset, the level of noise in quantum hardware, and the computational resources needed for implementation.[32][33] To address these uncertainties, this study conducts a comparative benchmarking of QSVM against multiple classical ML models across diverse datasets, including peptides and long protein sequences. This analysis is essential for understanding whether QSVM offers any real performance benefits, such as improved accuracy or computational efficiency, over traditional methods.[6] Additionally, it evaluates the computational overhead of QSVM, particularly in terms of training time and resource consumption to determine its feasibility for large-scale applications.This study also aims to determine if specific types of datasets are inherently more compatible with quantum-enhanced learning.[34] By examining these factors, this research provides a comprehensive evaluation of QSVM’s current capabilities, shedding light on its strengths and limitations in real-world applications.

### 1.4 Research Objective

The primary goal of this research is to systematically compare QSVM with classical machine learning models such as classical SVM across four datasets. This analysis examines critical performance indicators, including accuracy, precision, recall, and F1-score, to evaluate the performance of QSVM compared to traditional techniques.[35] Additionally, the study explores computational efficiency by analyzing training time, inference speed, and resource usage.[36] By doing this, it offers important insights into the practical feasibility of QSVM in real-world applications. Another key focus of this research is scalability.[37] As dataset size and feature complexity grow, it is important to determine whether QSVM maintains its performance advantages or runs into computational bottlenecks. Furthermore, this study explores the specific conditions where QSVM outperforms classical ML models and where it may still lag behind.[38] By exploring these trade-offs, we seek to offer a better understanding of the present challenges faced by quantum computing in machine learning applications.[39][40] Lastly, this research assesses the real-world practicability of QSVM, taking into account the existing constraints of running quantum models on classical hardware.[6] Through these objectives, this study aims to offer a comprehensive evaluation of QSVM’s applicabilities and its effectiveness compared to proven classical machine learning methods.

## 2. Related Works

### 2.1 Prior Research on Quantum SVM and Classical ML Benchmarking

Due to their great ability to define optimal decision boundaries in high-dimensional feature spaces, Support Vector Machines (SVMs) have become a mainstay in classification and regression tasks.[19][27] Quantum Support Vector Machines (QSVM) offer a new approach by utilizing quantum computing principles, including quantum kernel estimation and the mapping of data to high-dimensional quantum states are key concepts in this approach.[17][28] These advancements aim to enhance classical SVMs by potentially improving computational efficiency and performance in complex classification problems.[41] Several studies have explored the theoretical benefits of QSVM compared to its classical counterpart. One of the earliest and most influential works in this area was conducted by Rebentrost et al. (2014), who introduced a quantum algorithm for SVMs that demonstrated a possible exponential speedup in computing inner products for kernel methods.[42] Their research laid the groundwork for quantum-enhanced machine learning models, suggesting that QSVM could outperform classical SVMs in certain applications. Building on this foundation, Havlíček et al. (2019) explored Quantum Kernel Methods using variational quantum circuits and evaluated their performance on small datasets.[26] Their findings showed that QSVM can effectively approximate high-dimensional feature maps. However, they also emphasized that its real-world viability is heavily dependent on advancements in quantum hardware, particularly in reducing noise and increasing qubit stability.[43]

On the classical machine learning side, the performance of several models, such as SVM, Random Forest, XGBoost, and Deep Learning, has been assessed on a variety of datasets.[44] Research by Pedregosa et al. (2011), best known for their contributions to the scikit-learn framework, along with several other studies, has explored how factors such as hyperparameter tuning, dataset size, and feature selection impact the generalization ability and efficiency of classical SVMs.[45] These studies have provided valuable insights into optimizing SVM performance for a large range of applications, highlighting both its strengths and limitations in real-world scenarios.[46][47] Despite these developments, research directly comparing QSVM with various classical machine learning models on real-world datasets remains limited. This work seeks to fill that gap by providing an impartial comparison of QSVM with classical SVM and other models across four diverse datasets.

### 2.2 Key Contributions in quantum computing for machine learning

The rapid progress in quantum computing has sparked extensive research into its applications in machine learning. Several foundational studies have shaped this evolving field, offering both theoretical frameworks and practical insights into how quantum computing could enhance ML tasks.[48][6] These contributions have paved the way for exploring quantum algorithms that may outperform classical methods in specific scenarios. Biamonte et al. (2017) offered a comprehensive review of Quantum Machine Learning (QML), investigating the potential for quantum advancements to accelerate various machine learning tasks.[<u>49</u> Lloyd et al. (2014) proposed quantum algorithms for principal component analysis (PCA) and clustering, showing how quantum computing can improve data processing capabilities.[50] Lloyd and Weedbrook (2018) introduced Quantum Generative Adversarial Learning, a concept that leverages quantum computing for generative adversarial networks and explores how quantum systems could be leveraged for generative modeling tasks.[51] Schuld et al. (2019) examined quantum feature maps, showing how they enable more powerful kernel-based learning techniques, especially for classification.[52] Cerezo et al. (2021) explored variational quantum algorithms, emphasizing their importance in optimization and machine learning.[53] Meanwhile, Abbas et al. (2020) highlighted the advantages of quantum neural networks, showing that they offer greater expressibility and faster training compared to classical models, making them promising for specific tasks.[54] These foundational research studies establish the theoretical basis for Quantum Machine Learning (QML) and highlight the need for further empirical evaluations.[55] While research suggests that quantum computing could enhance ML performance, benchmarking in real-world scenarios is essential for evaluating the feasibility and constraints of quantum-enhanced machine learning models.[56]

### 2.3 Gaps in Current Literature

Despite the increasing interest in QML, several key gaps remain in the research landscape.[57] A major challenge is the absence of practical benchmarking. While many studies highlight the theoretical advantages of QSVM, few have systematically tested its performance against multiple classical models across diverse real-world datasets.[58][59] As a result, questions about the practical effectiveness of QSVM, its accuracy, and efficiency in real-world applications remain largely unanswered. Another significant issue is the scalability of QSVM.[60] Prior research has primarily tested QSVM on synthetic or small-scale datasets, making it unclear whether these quantum-enhanced models can handle larger, high-dimensional datasets efficiently. Given that many practical ML applications involve extensive datasets, understanding QSVM’s ability to scale is crucial for its adoption in real-world tasks.[42][61]

The computational cost and hardware constraints of QSVM remain largely unexamined.[15] While QSVM holds theoretical promise, key factors such as quantum decoherence and noise both of which can significantly affect accuracy and stability are often overlooked.[62] These issues underscore the necessity for additional experimental research to evaluate whether QSVM provides any advantage over classical models, given the limitations of current quantum technology.[63] By filling these gaps, this research seeks to provide a more complete understanding of QSVM’s practical applicability. Through rigorous benchmarking against classical ML techniques across multiple datasets, we assess its performance and feasibility within the limitations of today’s quantum hardware.

## 3. Methodology

### 3.1 Datasets

In this study, we benchmark a comparison of Quantum Support Vector Machines (QSVM) with classical machine learning models across various datasets for protein and peptide classification..[64] These datasets contain sequential biological data, where amino acid sequences are transformed into numerical feature vectors. Feature scaling techniques were applied during preprocessing depending on the original work done in the paper for fair evaluation.[65] The detailed information about datasets used in this study can be found below:

#### 3.1.1 CLBTope

The CLBTope dataset is designed for B-cell epitope classification, containing sequences sourced from antibody-antigen structure complexes and the Immune Epitope Database (IEDB).[66][67] The dataset includes a training set of 6,296 samples and a test set comprising 1,575 samples. These sequences capture key biochemical and structural properties of amino acids, enabling the identification of potential B-cell epitopes. The dataset includes both linear epitopes, which are contiguous stretches of amino acids, and conformational epitopes formed by amino acids that are spatially adjacent in the three-dimensional structure of a protein, even though they may be far apart in the sequence. By incorporating diverse epitope types, CLBTope provides a comprehensive dataset for studying epitope prediction in immunoinformatics.

#### 3.1.2 ExoProPred

The ExoProPred dataset is created for the purpose of classifying exosomal and non-exosomal proteins, consisting of sequences sourced from UniProt and the ExoPred dataset.[68][69] It comprises 2,831 exosomal protein sequences and an equal number of non-exosomal protein sequences, ensuring a balanced classification task. The dataset was curated using strict preprocessing criteria to enhance sequence quality and reduce redundancy. To maintain diversity and eliminate excessive sequence similarity, redundant sequences were eliminated using CD-HIT software, applying a similarity threshold of below 40%.[70] Additionally, sequences containing non-standard amino acids ("BJOUXZ") and those shorter than 55 or longer than 1,500 amino acids were excluded to ensure biologically relevant data.

#### 3.1.3 HemoPI

For this study, we used the HemoPI-1 dataset, which is designed for distinguishing between hemolytic and non-hemolytic peptides, tackling a significant challenge in therapeutic peptide development of hemotoxicity.[71] HemoPI-1 was constructed using 552 experimentally validated hemolytic peptides from the Hemolytik database, selected based on potency thresholds such as EC50 ≤ 100 µM and MHC ≤ 250 µg/ml.[72] The dataset was built with a set of 552 non-hemolytic peptides randomly chosen from the Swiss-Prot database while maintaining a similar length distribution.[73] Additionally, five alternative negative datasets (Random1–Random5) were generated to ensure unbiased classification.

#### 3.1.4 ToxinPred3

The ToxinPred3 dataset is specifically designed to classify toxic and non-toxic peptides, playing a crucial role in peptide-based drug development and toxicity prediction.[74] It was constructed by compiling 5,518 experimentally validated toxic peptides from various databases, including Conoserver[75], DRAMP[76], CAMPR3[77], dbAMP2.0[78], YADAMP[79], DBAASP-v3[80], and UniProt[68]. To ensure biological relevance, peptides longer than 35 residues or containing non-natural amino acids were removed. To ensure a balanced classification task, an equal set of 5,518 non-toxic peptides was selected randomly from the Swiss-Prot database, maintaining similar length and composition criteria as the toxic peptides.

### 3.2 Feature Generation

To extract numerical representations from amino acid sequences, we utilized Pfeature, a well-established bioinformatics software known for its robust feature extraction capabilities from peptides and proteins.[81] Protein sequences were transformed into numerical feature vectors to capture their biochemical characteristics, enabling more context information for effective machine learning applications. Unlike other datasets that primarily focus on sequence availability, our approach ensures comprehensive preprocessing to enhance data reliability. The 20-dimensional AAC based feature vectors derived from Pfeature served as inputs for both traditional machine learning models and quantum support vector machines (QSVM).

### 3.3 Normalization

While no additional feature transformation techniques were applied during preprocessing to preserve the raw numerical representations, Min-Max scaling was later employed to standardize input values into a range of [0, 1].[82] This transformation ensures that all input features contribute proportionally to the learning process, enhancing the model’s capacity to identify significant patterns. The Min-Max scaling process is mathematically represented as follows:

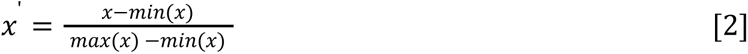

where x′ denotes the normalized value, x is the original feature value, and min(x) and max(x) represent the minimum and maximum values of the feature, respectively.

By standardizing the input data, we prevent biases caused by differences in scales. Thereby improving consistency in model training and generalization.[65]

### 3.4 Classical Machine Learning Models

To establish a solid benchmark, we compare Quantum Support Vector Machine (QSVM) with the widely recognized classical Support Vector Machine (SVM). SVM is an effective supervised learning algorithm that identifies the best decision boundary, or hyperplane, to distinguish data points within a high-dimensional space. SVM works by enlarging the gap between classes, ensuring a clearer separation, and enhances generalization reducing the risk of misclassification. To optimize SVM’s performance, we fine-tune key hyperparameters using grid search.[83][84] We went ahead with linear kernel type, allowing SVM to create complex decision boundaries by transforming input features into higher-dimensional spaces. The regularization parameter (C) manages the balance between maximizing the margin and reducing classification errors, aiding in the control of bias and variance.

### 3.5 Quantum SVM

QSVM is implemented using the PennyLane library, which provides a framework for quantum machine learning, leveraging automatic differentiation allows for smooth integration with quantum simulators and hardware.[85] The key steps in the QSVM implementation include quantum feature mapping, quantum kernel estimation, and classification using a quantum-enhanced SVM. In quantum feature mapping, classical data is transformed into a quantum state through parameterized quantum circuits, such as those using Pauli strings. This step is essential because it enables the quantum model to take advantage of high-dimensional Hilbert spaces for better separability.[86] In our study, we experimented with different numbers of qubits, including 6, 10, and 14, to examine the impact of the quantum feature space on classification performance. The quantum kernel trick is utilized by calculating the inner product of quantum state vectors, resulting in the creation of a quantum kernel matrix that replaces traditional kernel functions like RBF or polynomial. This kernel matrix is subsequently employed in an SVM classifier to distinguish data points within the quantum feature space. The quantum-enhanced SVM is trained on this kernel matrix using a classical optimization procedure. Due to current limitations in quantum hardware, experiments are primarily conducted using PennyLane’s quantum simulators but to assess its practical feasibility, quantum hardwares should be tested as well.

### 3.6 Cross Validation: Improving Generalization in Model Performance

For robust and unbiased evaluation, both classical and quantum models are trained and evaluated using stratified k-fold cross-validation.[87] This approach preserves the original class distribution across all folds, mitigating overfitting and enhancing the reliability of performance assessments. In our study, we employ k=5, partitioning the training data. The data is divided into five separate subsets (folds). In each iteration, k-1 folds are used for training, with the remaining fold acting as the validation set. This procedure is repeated across multiple iterations until each fold has been used for validation exactly once.[88] Notably, the testing set remains unchanged throughout the cross-validation process, ensuring a consistent basis for evaluation. After completing all iterations, performance metrics from each fold are compiled, and their mean value is computed. This averaging reduces variance and enhances the reliability of model comparisons. The results of various models are summarized in section 5.1.

### 4.6 Experimental Setup & Evaluation Metrics

The QSVM experiments were conducted using PennyLane’s quantum simulator within a Python 3.10 environment. To provide a thorough assessment, various classification metrics were measured, such as accuracy, precision, recall, specificity, F1-score, Matthews correlation coefficient (MCC), and the area under the receiver operating characteristic curve (AUC). These metrics offer a detailed evaluation of the models’ performance in distinguishing between classes.[89] The evaluation framework consists of both threshold-dependent and threshold-independent metrics. Accuracy, precision, recall (sensitivity), specificity, F1-score, and MCC are threshold-dependent metrics, as their values are influenced by the classification threshold. In contrast, AUC is a threshold-independent measure, it assesses the model’s capacity to differentiate between positive and negative classes at different decision thresholds. These well-established evaluation criteria ensure a robust and reliable benchmarking process. The mathematical expressions for the threshold-dependent metrics are outlined as follows:

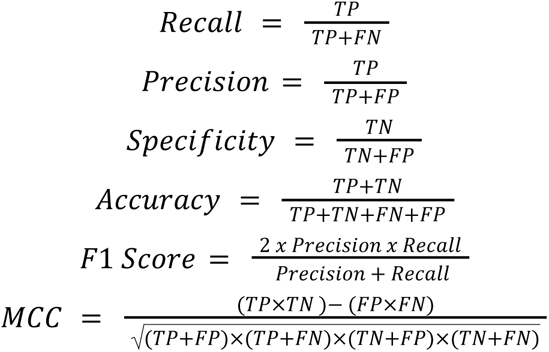

Here, FP, FN, TP, and TN represent false positives, false negatives, true positives, and true negatives, respectively.

Additionally, AUC is computed as a threshold-independent metric that measures the model’s ability to distinguish between positive and negative classes using the receiver operating characteristic (ROC) curve. Beyond classification performance, computational efficiency is also analyzed by comparing each model’s training and inference time. Furthermore, a scalability analysis is conducted to evaluate how performance varies with dataset size, including the number of qubits and feature complexity for quantum models and computational requirements for classical models.[35]

## 4. Results & Discussion

### 4.1 Comparative Analysis of Performance

In order to find the most effective predictive models for our classification tasks, we initiated our analysis by constructing composition-based features were extracted using the Pfeature software, specifically utilizing Amino Acid Composition (AAC) [81], which captures the frequency of each amino acid in a sequence, AAC provides a fixed-length numerical representation that is ideal for subsequent machine learning tasks. This feature engineering technique was applied for all four benchmark datasets - CLBTope, ExoProPred, Hemo-Pi, and ToxinPred3, which served as the standardized input for both classical machine learning (CML) models and Quantum Support Vector Machine (QSVM), a quantum-enhanced variant of the classical SVM.

A diverse set of CML models was trained and evaluated, including Extra Trees (ET), Support Vector Classifier (SVC), Random Forest (RF), Logistic Regression (Logit) and Extreme Gradient Boosting (XGB), reported along with their baseline models from prior studies. These ensemble and kernel-based models are widely recognized for their robustness, predictive power, and capacity to handle high-dimensional data. Each model evaluation was conducted using a standard set of performance metrics, such as sensitivity, specificity, accuracy, precision, F1-score, Matthews Correlation Coefficient (MCC), and Area Under the ROC Curve (AUROC). Both training and validation results were recorded to assess each model’s ability to generalize beyond the training data and recognize potential overfitting. To provide a fair comparison, the QSVM was benchmarked using the same AAC-based features, ensuring consistency in input representation across all models. To evaluate the influence of quantum computational resources, QSVM was trained using three different qubit configurations - 6, 10, and 14 qubits. This provides an empirical analysis of how the number of quantum features impacts model performance, generalization, and convergence behavior. The comprehensive outcomes of these experiments are in the following tables.

For the CLBTope dataset, Extra Trees emerged as the most robust classifier, achieving the highest validation metrics with an AUROC of 0.82 and MCC of 0.49. In contrast, QSVM showing improved performance with more qubits, only reached AUROC of 0.68 and MCC of 0.37 at 14 qubits, indicating that although quantum models have potential, classical tree-based ensembles remain superior in B-cell epitope classification.

On the ExoProPred dataset, performance differences between QSVM and classical models were narrower. The best classical models, such as SVC and XGB, achieved validation accuracy around 0.66, MCC ∼0.32, and AUROC ∼0.72. QSVM showed comparable accuracy (0.66) and MCC (0.32) at 14 qubits, but slightly lagged in AUROC (0.66), indicating reasonable generalization power but slightly weaker discrimination of positive and negative classes.

The Hemo-Pi dataset showed the strongest QSVM performance across all datasets. At 14 qubits, QSVM achieved a validation accuracy of 0.95, an F1-score of 0.93, and an MCC of 0.91, closely matching or even slightly exceeding classical models such as SVM and Logistic Regression. The classical SVM achieved similar accuracy (0.95), but lower F1-score (0.92) and MCC (0.83), suggesting that QSVM generalizes better in this biologically complex task, possibly due to its ability to capture subtle patterns in the feature space.

For the ToxinPred3 dataset, classical models clearly outperformed QSVM. The Extra Trees classifier achieved an AUROC of 0.94 and an MCC of 0.75 on the validation set significantly surpassing QSVM’s best performance at 14 qubits which is an AUROC of 0.84 and an MCC of 0.68. Other classical models, such as RF and XGB also maintained consistent performance advantages over QSVM in this task.

In summary, while QSVM demonstrates competitive and promising results, especially at higher qubit counts, it does not consistently outperform classical models across all datasets. Its effectiveness appears to be highly dataset-dependent, with the best results observed in the Hemo-Pi dataset. The performance improvements with increased qubits suggest that richer quantum kernels can better capture sequence patterns. However, the superior and stable performance of classical models across various datasets highlights the current limitations of quantum machine learning. These results highlight the importance of continued research in quantum kernel optimization and hardware scalability to enhance QSVM’s practical utility in real-world bioinformatics scenarios.

### 4.2 Computational Efficiency

The computational performance of the Quantum Support Vector Machine (QSVM) was analyzed by measuring its training and inference times across four benchmark datasets. In comparison, classical machine learning (CML) models, especially tree-based methods like Random Forest (RF) and Extra Trees (ET) are also tested. We know that CMLs are well-optimized and benefit from hardware acceleration, including GPU support, enabling them to train and infer rapidly with minimal overhead.[90] QSVM, on the other hand, introduces significant computational challenges. The primary sources of this overhead are quantum kernel estimation and quantum circuit execution, both of which are computationally intensive, especially when simulated on classical hardware.[29] These challenges are exacerbated in simulated quantum environments, where additional latency is introduced, and on actual quantum hardware, where noise and limited qubit connectivity further impact performance.[91] The computational time analysis for QSVM on classical hardware is observed in the below figure.

Figure 1 illustrates that, while increasing the number of qubits from 6 to 14 leads to marginal scalability improvements, the overall computational cost remains substantial. This is largely because of the exponential increase in the number of parameters and circuit complexity with more qubits, making real-time or large-scale deployment of QSVM currently impractical. Thus, although QSVM exhibits potential, its computational feasibility remains constrained.[92] The trade-off between model accuracy and computational efficiency indicates that, under current technological limitations, QSVM is not yet competitive with classical models in terms of performance.[62]

**Figure 1:**
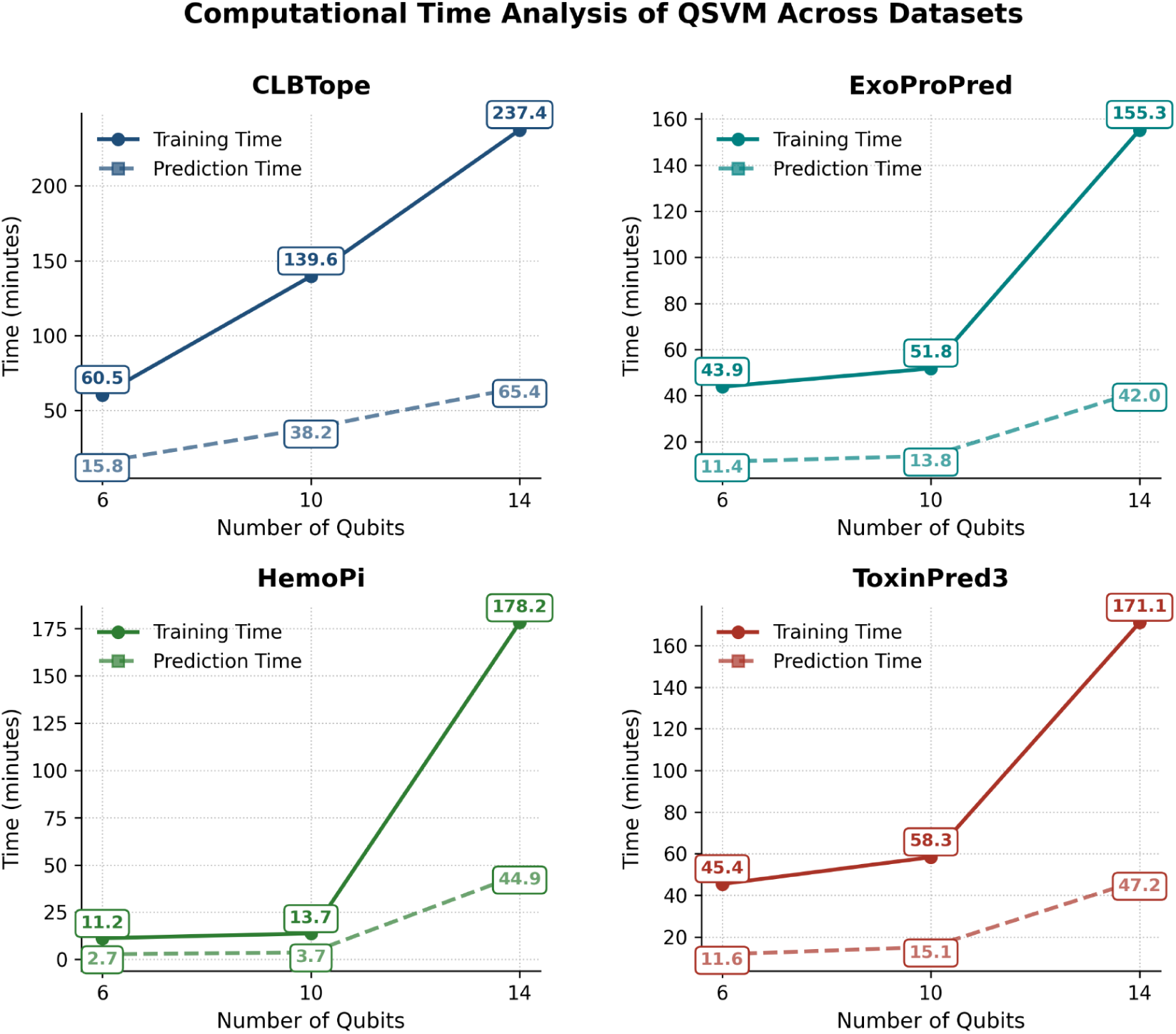
Training and Inference Time of QSVM Across Different Qubit Counts.

To contextualize these findings, we compared the computational times of QSVM with those of classical models, displayed in the figure below.

As shown in Figure 2, CML models consistently outperformed QSVM in terms of training and inference speed. This efficiency gap is especially evident in datasets such as Hemo-Pi, where classical models completed training and prediction within seconds, while QSVM took significantly longer, reinforcing the disparity in computational efficiency. Similar trends were observed across other datasets, where CML’s optimized algorithms enabled fast training and inference, while QSVM exhibited significant latency due to quantum processing. Further emphasizing the resource-intensive nature of quantum models.

**Figure 2:**
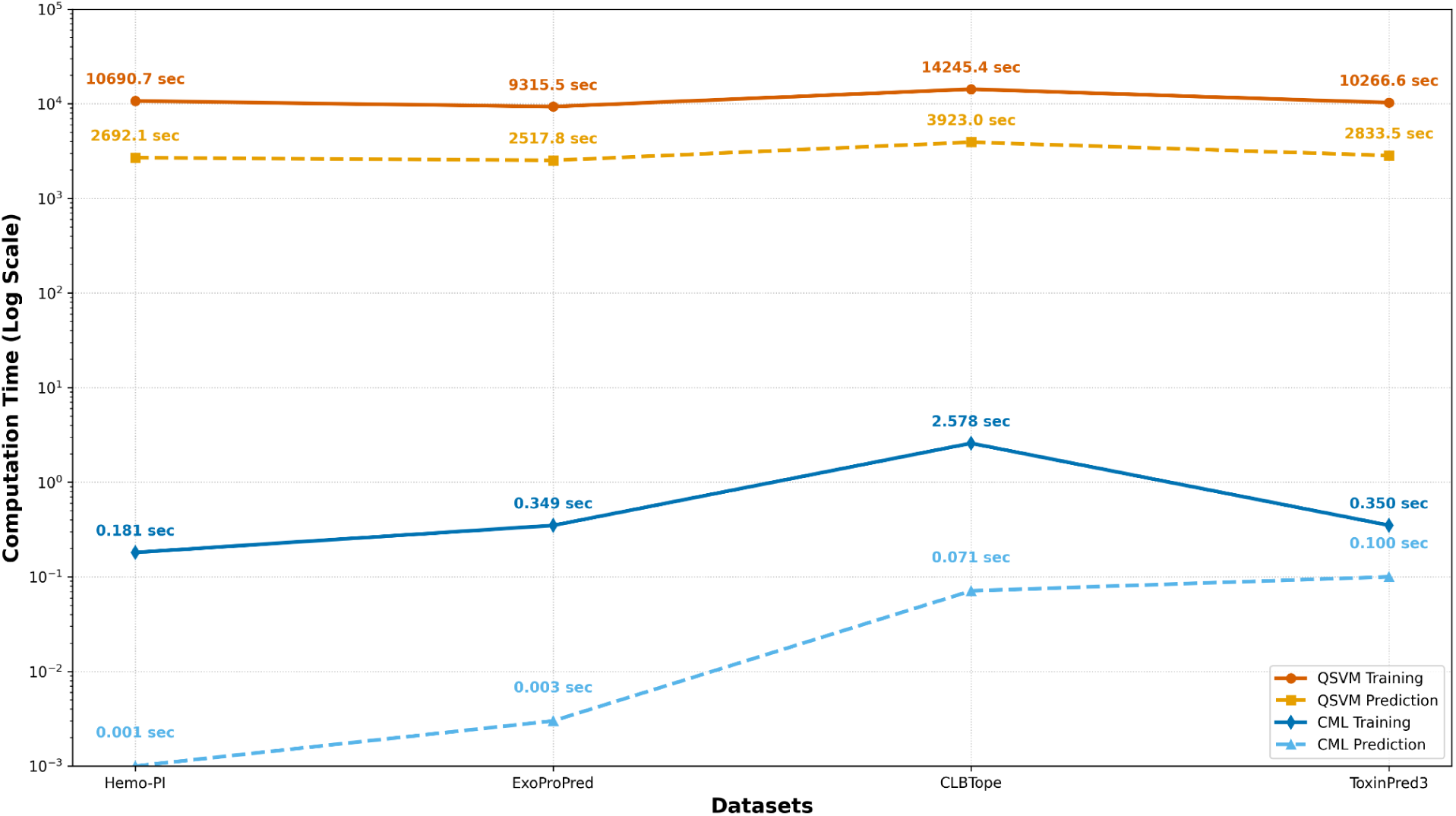
Computational Training and Inference Time of QSVM and CML.

These results highlight the current limitations of QSVM in terms of computational feasibility. Despite its theoretical promise, QSVM’s heavy resource requirements and execution latency present significant barriers to practical adoption. Tree-based classical models, by contrast, offer both speed and accuracy, making them far more appropriate for real-time applications and extensive datasets under present hardware capabilities.

### 4.3 Overall Summary

To evaluate the scalability of the Quantum Support Vector Machine (QSVM), we analyzed its classification performance across increasing qubit configurations, with highest scores observed at the 14-qubit level. Table 5 summarizes the Area Under the Curve (AUC) scores for both QSVM and CML models across four therapeutic sequential datasets, offering a direct performance comparison between quantum and classical approaches. Comprehensive results for all qubit configurations are provided in the supplementary materials.

**Table 1:** Performance of composition-based features of CLBTope dataset.

| Models |  | Training |  |  |  |  |  |  | Validation |  |  |  |  |  |  |
| --- | --- | --- | --- | --- | --- | --- | --- | --- | --- | --- | --- | --- | --- | --- | --- |
|  |  | Sens | Spec | Acc | Prec | F1 | MCC | AUC | Sens | Spec | Acc | Prec | F1 | MCC | AUC |
| Baseline | RF | 0.71 | 0.70 | 0.71 | - | - | 0.42 | **0.79** | 0.73 | 0.72 | 0.72 | - | - | 0.45 | **0.79** |
| High-Scoring | ET | 0.72 | 0.72 | 0.72 | 0.71 | 0.72 | 0.44 | **0.79** | 0.74 | 0.75 | 0.74 | 0.74 | 0.74 | 0.49 | **0.82** |
|  | XGB | 0.69 | 0.71 | 0.70 | 0.70 | 0.70 | 0.41 | **0.78** | 0.71 | 0.73 | 0.72 | 0.72 | 0.71 | 0.44 | **0.80** |
|  | RF | 0.66 | 0.71 | 0.68 | 0.68 | 0.67 | 0.37 | **0.75** | 0.67 | 0.73 | 0.70 | 0.70 | 0.68 | 0.40 | **0.77** |
| QSVM with various qubits | 6 | 0.52 | 0.56 | 0.54 | 0.53 | 0.53 | 0.08 | **0.54** | 0.49 | 0.59 | 0.54 | 0.54 | 0.52 | 0.08 | **0.54** |
|  | 10 | 0.59 | 0.67 | 0.63 | 0.63 | 0.61 | 0.25 | **0.63** | 0.65 | 0.63 | 0.64 | 0.63 | 0.64 | 0.28 | **0.64** |
|  | 14 | 0.65 | 0.66 | 0.66 | 0.65 | 0.65 | 0.31 | **0.66** | 0.69 | 0.68 | 0.69 | 0.67 | 0.68 | 0.37 | **0.68** |

**Table 2:** Performance of composition-based features of ExoProPred dataset.

| Models |  | Training |  |  |  |  |  |  | Validation |  |  |  |  |  |  |
| --- | --- | --- | --- | --- | --- | --- | --- | --- | --- | --- | --- | --- | --- | --- | --- |
|  |  | Sens | Spec | Acc | Prec | F1 | MCC | AUC | Sens | Spec | Acc | Prec | F1 | MCC | AUC |
| Baseline | SVC | 0.65 | 0.65 | 0.65 | - | - | 0.30 | **0.71** | 0.66 | 0.66 | 0.66 | - | - | 0.32 | **0.71** |
| High-Scoring | XGB | 0.66 | 0.63 | 0.64 | 0.64 | 0.65 | 0.29 | **0.70** | 0.67 | 0.64 | 0.66 | 0.65 | 0.66 | 0.32 | **0.72** |
|  | SVC | 0.66 | 0.61 | 0.64 | 0.63 | 0.65 | 0.28 | **0.70** | 0.65 | 0.64 | 0.65 | 0.64 | 0.65 | 0.29 | **0.70** |
|  | RF | 0.65 | 0.61 | 0.63 | 0.62 | 0.64 | 0.26 | **0.69** | 0.67 | 0.61 | 0.64 | 0.63 | 0.65 | 0.28 | **0.70** |
| QSVM with various qubits | 6 | 0.65 | 0.63 | 0.64 | 0.63 | 0.64 | 0.27 | **0.64** | 0.64 | 0.66 | 0.65 | 0.65 | 0.65 | 0.29 | **0.65** |
|  | 10 | 0.66 | 0.64 | 0.65 | 0.64 | 0.65 | 0.29 | **0.64** | 0.66 | 0.67 | 0.66 | 0.66 | 0.66 | 0.32 | **0.66** |
|  | 14 | 0.66 | 0.63 | 0.64 | 0.64 | 0.65 | 0.29 | **0.64** | 0.66 | 0.67 | 0.66 | 0.66 | 0.66 | 0.32 | **0.66** |

**Table 3:** Performance of composition based features of Hemo-Pi dataset.

| Models |  | Training |  |  |  |  |  |  | Validation |  |  |  |  |  |  |
| --- | --- | --- | --- | --- | --- | --- | --- | --- | --- | --- | --- | --- | --- | --- | --- |
|  |  | Sens | Spec | Acc | Prec | F1 | MCC | AUC | Sens | Spec | Acc | Prec | F1 | MCC | AUC |
| Baseline | SVM | 0.95 | 0.95 | 0.95 | - | - | 0.91 | - | 0.96 | 0.99 | 0.96 | - | - | 0.93 | - |
| High-Scoring | SVM | 0.95 | 0.93 | 0.94 | 0.94 | 0.94 | 0.88 | **0.98** | 0.95 | 0.87 | 0.91 | 0.88 | 0.92 | 0.83 | **0.98** |
|  | Logit | 0.94 | 0.93 | 0.94 | 0.94 | 0.94 | 0.88 | **0.98** | 0.95 | 0.89 | 0.92 | 0.90 | 0.93 | 0.85 | **0.98** |
|  | RF | 0.92 | 0.92 | 0.92 | 0.92 | 0.92 | 0.84 | **0.97** | 0.95 | 0.93 | 0.94 | 0.93 | 0.94 | 0.88 | **0.98** |
| QSVM with various qubits | 6 | 0.72 | 0.85 | 0.78 | 0.83 | 0.76 | 0.57 | **0.78** | 0.71 | 0.85 | 0.78 | 0.83 | 0.76 | 0.57 | **0.78** |
|  | 10 | 0.77 | 0.97 | 0.87 | 0.96 | 0.86 | 0.75 | **0.87** | 0.80 | 0.98 | 0.89 | 0.97 | 0.88 | 0.79 | **0.89** |
|  | 14 | 0.94 | 0.96 | 0.95 | 0.96 | 0.95 | 0.90 | **0.95** | 0.98 | 0.93 | 0.95 | 0.93 | 0.96 | 0.91 | **0.95** |

**Table 4:** Performance of composition-based features of Toxinpred3 dataset.

| Models |  | Training |  |  |  |  |  |  | Validation |  |  |  |  |  |  |
| --- | --- | --- | --- | --- | --- | --- | --- | --- | --- | --- | --- | --- | --- | --- | --- |
|  |  | Sens | Spec | Acc | Prec | F1 | MCC | AUC | Sens | Spec | Acc | Prec | F1 | MCC | AUC |
| Baseline | ET | 0.83 | 0.91 | 0.87 | 0.90 | 0.86 | 0.74 | **0.94** | 0.85 | 0.90 | 0.87 | 0.89 | 0.87 | 0.75 | **0.94** |
| High-Scoring | ET | 0.87 | 0.85 | 0.86 | 0.85 | 0.86 | 0.73 | **0.93** | 0.89 | 0.84 | 0.86 | 0.85 | 0.87 | 0.73 | **0.93** |
|  | XGB | 0.83 | 0.87 | 0.85 | 0.87 | 0.85 | 0.71 | **0.93** | 0.85 | 0.86 | 0.86 | 0.86 | 0.86 | 0.71 | **0.93** |
|  | RF | 0.82 | 0.9 | 0.86 | 0.89 | 0.86 | 0.73 | **0.93** | 0.84 | 0.89 | 0.86 | 0.88 | 0.86 | 0.72 | **0.93** |
| QSVM<br>with<br>various<br>qubits | 6 | 0.84 | 0.88 | 0.86 | 0.88 | 0.86 | 0.73 | 0.86 | 0.81 | 0.85 | 0.83 | 0.84 | 0.82 | 0.66 | 0.83 |
|  | 10 | 0.87 | 0.90 | 0.88 | 0.89 | 0.88 | 0.77 | 0.88 | 0.83 | 0.85 | 0.84 | 0.85 | 0.84 | 0.69 | 0.84 |
|  | 14 | 0.87 | 0.90 | 0.88 | 0.89 | 0.88 | 0.77 | **0.88** | 0.83 | 0.84 | 0.84 | 0.84 | 0.83 | 0.68 | **0.84** |

**Table 5:**
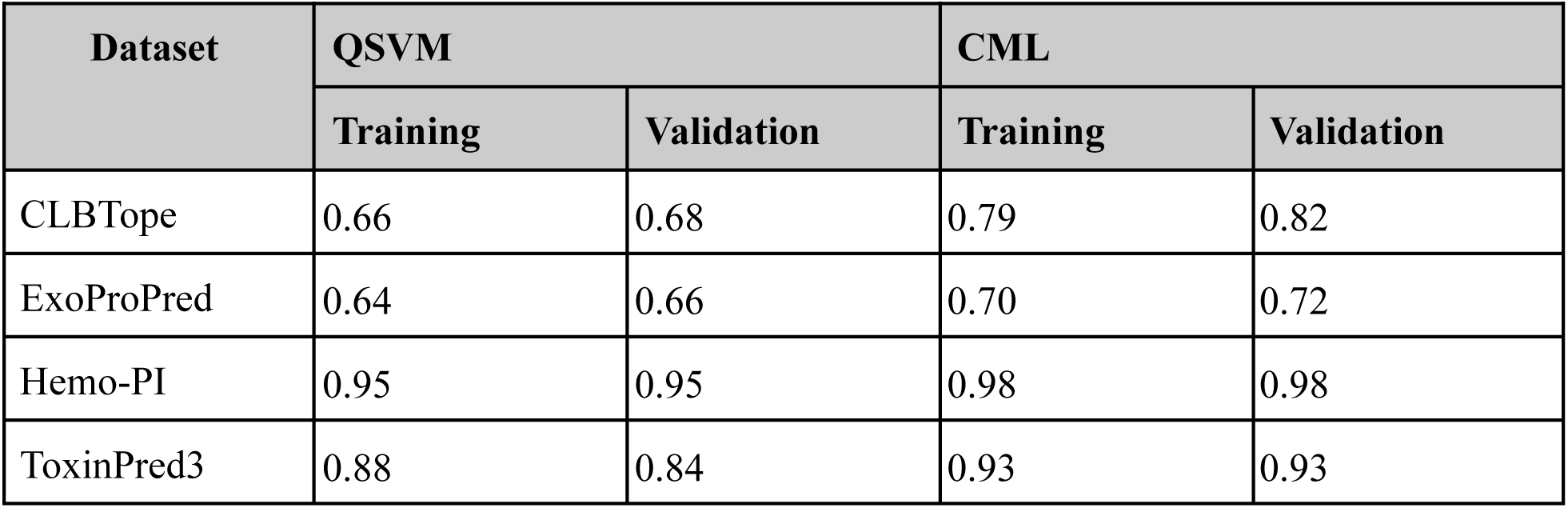
Performance Comparison of QSVM and CML Models based on Area Under the Curve.

As observed in Table 5, the overall QSVM performance improved with an increased number of qubits, supporting the idea that higher-dimensional quantum feature maps enhance the model’s capacity to learn complex patterns. However, this improvement tends to plateau beyond a certain qubit threshold, indicating diminishing returns with added quantum resources. For instance, in the Hemo-PI dataset, QSVM achieved strong results with relatively fewer qubits, suggesting its potential for low-resource quantum applications. On the other hand, datasets like CLBTope and ToxinPred3 demonstrated limited gains, highlighting the dependency of QSVM’s effectiveness on data characteristics.

Despite these promising results, classical models, particularly tree-based models and support vector machines consistently outperformed QSVM in both training and validation across most datasets. These models offer improved computational efficiency and scalability, making them better suited for practical deployment in real-world scenarios under current hardware limitations.

QSVM introduces a novel approach via quantum kernel estimation, offering a fresh perspective on feature representation in high-dimensional spaces. However, its reliance on complex quantum circuitry, increased computation time, and sensitivity to hardware noise limits its immediate practicality. These constraints emphasize the necessity for enhanced quantum hardware and error-mitigation strategies to fully realize its potential.

In summary, while QSVM demonstrates promising potential in specific contexts, it remains outperformed by classical machine learning models in terms of both accuracy and computational efficiency. This study highlights both the strengths and limitations of QSVM, like introducing innovative quantum kernel methods while also revealing current limitations related to scalability and practical deployment. Classical models continue to dominate in large-scale, real-world settings due to their proven robustness and efficiency. Moving forward, hybrid quantum-classical approaches that integrate the expressive capabilities of quantum kernels with the reliability of classical algorithms may help close the performance gap and pave the way for the practical adoption of quantum-enhanced machine learning systems.

## 5. Conclusion & Implications for Future Work

This study benchmarked Quantum Support Vector Machines (QSVM) were compared with classical machine learning (CML) models across four therapeutic protein and peptide datasets. While QSVM introduces a novel paradigm through quantum kernel estimation and demonstrates performance gains with increased qubit counts, it consistently lags behind classical models such as Random Forest, Extra Trees, and Support Vector Machines in terms of accuracy, computational efficiency, and scalability.

Although quantum techniques hold promise across many scientific domains, our results reaffirm that classical models remain more dependable for real-world classification tasks, particularly under current hardware limitations.[93] QSVM’s practical limitations stem from quantum-specific challenges, including circuit noise, decoherence, limited qubit availability, and the computational overhead of quantum kernel evaluations.[94][95] Moreover, the effectiveness of QSVM is highly dataset-dependent, with its advantages only emerging under specific data structures and encoding strategies.

Looking ahead, the path forward lies in hybrid quantum-classical approaches like integrating quantum kernels with classical algorithms to balance expressiveness and efficiency. Future work should explore adaptive quantum encoding methods tailored to specific types of data, improvements in quantum circuit design, and error-resilient computation. As quantum hardware evolves with better error correction, qubit fidelity, and scalability, QSVM may emerge as a more practical alternative.

In conclusion, while QSVM offers a compelling direction for computational biology and quantum machine learning, it remains an experimental approach at present as its real-world applicability remains constrained by computational limitations and dataset dependencies. Continued innovation in hybrid frameworks, hardware development, and algorithmic optimizations will be essential to unlocking its full potential for large-scale applications and beyond.

## Authors’ contributions

GPSR collected the datasets. PT processed the datasets and developed the prediction models. PT and NKM implemented the algorithms. GPSR analyzed the results, and PT performed the benchmarking. PT and GPSR penned the manuscript. GPSR conceived and coordinated the project. All authors have read and approved the final manuscript.

## Acknowledgment

Authors are thankful to the University Grants Commission (UGC), and Department of Biotechnology (DBT) for fellowships and financial support, and the Department of Computational Biology, IIITD New Delhi for infrastructure and facilities.

## Abbreviations

QML: Quantum Machine Learning
CML: Classical Machine Learning
VQC: Variational Quantum Circuits
QSVM: Quantum Support Vector Machines
QNN: Quantum Neural Networks
PCA: Principal Component Analysis
MCC: Matthews Correlation Coefficient
AUC: Area under the receiver operating characteristic curve
RF: Random Forest
ET: Extra Trees
PCC: Pearson correlation coefficient
R^2^: Coefficient of determination

## References

1. Goyal, R.K., Maharaj, S., Kumar, P. et al. Exploring quantum materials and applications: a review. J Mater. Sci: Mater Eng. 20, 4 (2025). 10.1186/s40712-024-00202-7

2. Memon, Q. A., Al Ahmad, M., & Pecht, M. (2024). Quantum Computing: Navigating the Future of Computation, Challenges, and Technological Breakthroughs. Quantum Reports, 6(4), 627–663. 10.3390/quantum6040039

3. Qi, J., Yang, C., Chen, S. Y., & Chen, P. (2024, November 14). Quantum Machine Learning: An interplay between quantum computing and machine learning. arXiv.org. https://arxiv.org/abs/2411.09403

4. Du, Y., Wang, X., Guo, N., Yu, Z., Qian, Y., Zhang, K., Hsieh, M., Rebentrost, P., & Tao, D. (2025, February 3). Quantum Machine Learning: A hands-on tutorial for machine learning practitioners and researchers. arXiv.org. https://arxiv.org/abs/2502.01146

5. Samuel Yen-Chi Chen and Joongheon Kim. 2024. Hands-On Introduction to Quantum Machine Learning. In Proceedings of the 33rd ACM International Conference on Information and Knowledge Management (CIKM ’24). Association for Computing Machinery, New York, NY, USA, *5507–*5510. 10.1145/3627673.3679103

6. Lu, W., Lu, Y., Li, J., Sigov, A., Ratkin, L., & Ivanov, L. A. (2024). Quantum machine learning: Classifications, challenges, and solutions. Journal of Industrial Information Integration, 42, 100736. 10.1016/j.jii.2024.100736

7. Pugliese, R., Regondi, S., & Marini, R. (2021). Machine learning-based approach: global trends, research directions, and regulatory standpoints. Data Science and Management, 4, 19–29. 10.1016/j.dsm.2021.12.002

8. Rashid, A. B., & Kausik, M. a. K. (2024). AI Revolutionizing Industries Worldwide: A comprehensive overview of its diverse applications. Hybrid Advances, 7, 100277. 10.1016/j.hybadv.2024.100277

9. Sheth, V., Tripathi, U., & Sharma, A. (2022). A Comparative analysis of Machine learning Algorithms for classification purpose. Procedia Computer Science, 215, 422–431. 10.1016/j.procs.2022.12.044

10. Cho, H., & Lee, S. (2021). Data Quality Measures and Efficient Evaluation Algorithms for Large-Scale High-Dimensional Data. Applied Sciences, 11(2), 472. 10.3390/app11020472

11. L’Heureux, A., Grolinger, K., Elyamany, H. F., & Capretz, M. A. M. (2017). Machine learning with big data: Challenges and approaches. IEEE Access, 5, 7776–7797. 10.1109/ACCESS.2017.2696365

12. Karim, M. R., et al. (2023). Interpreting black-box machine learning models for high-dimensional datasets. 2023 IEEE 10th International Conference on Data Science and Advanced Analytics (DSAA), 1–10. 10.1109/DSAA60987.2023.10302562

13. Tomar, S., Tripathi, R., & Kumar, S. (2025, January 16). Comprehensive Survey of QML: From data analysis to Algorithmic Advancements. arXiv.org. https://arxiv.org/abs/2501.09528

14. de Leon, N. P., et al. (2021). Materials challenges and opportunities for quantum computing hardware. Science, 372, eabb2823. 10.1126/science.abb2823

15. Fellous-Asiani, M., Chai, J. H., Whitney, R. S., & Auffèves, A., & Ng, H. K. (2021). Limitations in quantum computing from resource constraints. PRX Quantum, 2(4), 040335. 10.1103/PRXQuantum.2.040335

16. Beatrice, B. A., & Selvam, A. S. R. (2024). Quantum speedup in machine learning: Applications & challenges. International Journal of Research Publication and Reviews, 5(12), 1325–1326. https://ijrpr.com/uploads/V5ISSUE12/IJRPR36197.pdf

17. Yin, T. (2024). Quantum support vector machines: Theory and applications. Theoretical and Natural Science, 51, 34–42. 10.54254/2753-8818/51/2024CH015

18. McCarroll, R. (2024, August). *The future of machine learning: Expert predictions validated by AI research.* ResearchGate. https://www.researchgate.net/publication/383181593

19. Hearst, M. A., Dumais, S. T., Osuna, E., Platt, J., & Scholkopf, B. (1998). Support vector machines. IEEE Intelligent Systems and Their Applications, 13(4), 18–28. 10.1109/5254.708428

20. 20. Fürnkranz, J. (2011). Decision tree. In C. Sammut & G. I. Webb (Eds.), Encyclopedia of machine learning. Springer. 10.1007/978-0-387-30164-8_204

21. 21. Macukow, B. (2016). Neural networks – State of the art, brief history, basic models, and architecture. In 15th IFIP International Conference on Computer Information Systems and Industrial Management (CISIM) (pp. 3–14). Vilnius, Lithuania. 10.1007/978-3-319-45378-1_1

22. Mienye, I. D., & Sun, Y. (2022). A survey of ensemble learning: Concepts, algorithms, applications, and prospects. IEEE Access, 10, 99129–99149. 10.1109/ACCESS.2022.3207287

23. 23. Keogh, E., & Mueen, A. (2017). Curse of dimensionality. In C. Sammut & G. I. Webb (Eds.), Encyclopedia of machine learning and data mining. Springer. 10.1007/978-1-4899-7687-1_192

24. S. Misra and P. Rani, "Quantum Machine Learning: A Comprehensive Overview and Analysis," 2024 15th International Conference on Computing Communication and Networking Technologies (ICCCNT), Kamand, India, 2024, pp. 1-5, 10.1109/ICCCNT61001.2024.10724803

25. Zaman, K., Marchisio, A., Hanif, M. A., & Shafique, M. (2023). A survey on quantum machine learning: Current trends, challenges, opportunities, and the road ahead. arXiv preprint arXiv:2310.10315. https://arxiv.org/abs/2310.10315

26. Havlíček, V., Córcoles, A. D., Temme, K., Harrow, A. W., Kandala, A., Chow, J. M., & Gambetta, J. M. (2019). Supervised learning with quantum-enhanced feature spaces. Nature, 567(7747), 209–212. 10.1038/s41586-019-0980-2

27. Cervantes, J., Garcia-Lamont, F., Rodríguez-Mazahua, L., & Lopez, A. (2020). A comprehensive survey on support vector machine classification: Applications, challenges, and trends. Neurocomputing, 408, 189–215. 10.1016/j.neucom.2019.10.118

28. Zhou, X., Yu, J., Tan, J., &, et al. (2024). Quantum kernel estimation-based quantum support vector regression. Quantum Information Processing, 23, 29. 10.1007/s11128-023-04231-7

29. Innan, N., Khan, M., Panda, B., & Bennai, M. (2023). Enhancing quantum support vector machines through variational kernel training. Quantum Information Processing, 22(10). 10.1007/s11128-023-04138-3

30. 30. Park, J., Quanz, B., Wood, S., Higgins, H., & Harishankar, R. (2020, December 14). Practical application improvement to Quantum SVM: theory to practice. arXiv.org. https://arxiv.org/abs/2012.07725

31. Houssein, E. H., Abohashima, Z., Elhoseny, M., & Mohamed, W. M. (2022). Machine learning in the quantum realm: The state-of-the-art, challenges, and future vision. Expert Systems with Applications, 194, 116512. 10.1016/j.eswa.2022.116512

32. B.H., D. M., Pulicherla, P., M., P., & P, N. (2024). Quantum machine learning: bridging the GAP between classical and quantum computing. ITEGAM-JETIA, 10(48), 122–128. 10.5935/jetia.v10i48.943

33. Schetakis, N., Aghamalyan, D., Griffin, P., & Boguslavsky, M. (2022). Review of some existing QML frameworks and novel hybrid classical-quantum neural networks realising binary classification for the noisy datasets. Scientific reports, 12(1), 11927. 10.1038/s41598-022-14876-6

34. Ranga, D., Rana, A., Prajapat, S., Kumar, P., Kumar, K., & Vasilakos, A. V. (2024). Quantum Machine Learning: Exploring the Role of Data Encoding Techniques, Challenges, and Future Directions. Mathematics, 12(21), 3318. 10.3390/math12213318

35. Ajibosin, S. S., & Cetinkaya, D. (2024). Implementation and Performance Evaluation of Quantum Machine Learning Algorithms for Binary Classification. Software, 3(4), 498–513. 10.3390/software3040024

36. Preuveneers, D., Tsingenopoulos, I., & Joosen, W. (2020). Resource Usage and Performance Trade-offs for Machine Learning Models in Smart Environments. *Sensors (Basel*, Switzerland), 20(4), 1176. 10.3390/s20041176

37. Wang, J., & Boukerche, A. (2020). The scalability analysis of machine learning-based models in road traffic flow prediction. ICC 2020 - 2020 IEEE International Conference on Communications (ICC), 1–6. 10.1109/ICC40277.2020.9148964

38. Genç, S. (2024). Performance analysis of quantum and classical machine learning models for feature selection and classification of the diabetes health indicators dataset. 2024 8th International Artificial Intelligence and Data Processing Symposium (IDAP), 1–7. 10.1109/IDAP64064.2024.10710904

39. Nguyen, T., Sipola, T., & Hautamäki, J. (2024). Machine Learning Applications of Quantum Computing: A review. European Conference on Cyber Warfare and Security, 23(1), 322–330. 10.34190/eccws.23.1.2258

40. Peral-García, D., Cruz-Benito, J., & García-Peñalvo, F. J. (2024). Systematic literature review: Quantum machine learning and its applications. Computer Science Review, 51, 100619. 10.1016/j.cosrev.2024.100619

41. Anguita, D., Ridella, S., Rivieccio, F., & Zunino, R. (2003). Quantum optimization for training support vector machines. Neural Networks, 16(5–6), 763–770. 10.1016/s0893-6080(03)00087-x

42. Rebentrost, P., Mohseni, M., & Lloyd, S. (2014). Quantum support vector machine for big data classification. Physical Review Letters, 113(13), 130503. 10.1103/PhysRevLett.113.130503

43. You, X., Huang, Z., Alyanak, U., Romanenko, A., Grassellino, A., & Zhu, S. (2022). Stabilizing and improving qubit coherence by engineering the noise spectrum of two-level systems. Physical Review Applied, 18(4), 044026. 10.1103/PhysRevApplied.18.044026

44. Gupta, S., Saluja, K., Goyal, A., Vajpayee, A., & Tiwari, V. (2022). Comparing the performance of machine learning algorithms using estimated accuracy. Measurement: Sensors, 24, 100432. 10.1016/j.measen.2022.100432

45. Pedregosa, F., Varoquaux, G., Gramfort, A., Michel, V., Thirion, B., Grisel, O., Blondel, M., Prettenhofer, P., Weiss, R., Dubourg, V., Vanderplas, J., Passos, A., Cournapeau, D., Brucher, M., Perrot, M., & Duchesnay, É. (2011). Scikit-learn: Machine learning in Python. Journal of Machine Learning Research, 12, 2825–2830. https://dl.acm.org/doi/10.5555/1953048.2078195

46. 46. Kalita, D. J., Singh, V. P., & Kumar, V. (2020). A survey on SVM hyper-parameters optimization techniques. Retrieved from https://api.semanticscholar.org/CorpusID:215860154

47. Anguita, D., Ridella, S., Rivieccio, F., & Zunino, R. (2002). Automatic hyperparameter tuning for support vector machines. In J. R. Dorronsoro (Ed.), Artificial Neural Networks — ICANN 2002 (Vol. 2415). Springer, Berlin, Heidelberg. 10.1007/3-540-46084-5_217

48. Attya, S. M., Haddad, S. Q. G., Al-Zaidi, H. K. R., Hameed, W. M., & Latif, N. (2024). Quantum computing impact on traditional computer architecture models. Radioelectronics Nanosystems Information Technologies, 16(5), 691–704. 10.17725/j.rensit.2024.16.691

49. Biamonte, J., Wittek, P., Pancotti, N., Rebentrost, P., Wiebe, N., & Lloyd, S. (2017). Quantum machine learning. Nature, 549(7671), 195–202. 10.1038/nature23474

50. Lloyd, S., Mohseni, M., & Rebentrost, P. (2014). Quantum principal component analysis. Nature Physics, 10(9), 631–633. 10.1038/nphys3029

51. Lloyd, S., & Weedbrook, C. (2018). Quantum generative adversarial learning. Physical Review Letters, 121(4), 040502. 10.1103/PhysRevLett.121.040502

52. Schuld, M., & Killoran, N. (2019). Quantum machine learning in feature Hilbert spaces. Physical Review Letters, 122(4), 040504. 10.1103/PhysRevLett.122.040504

53. Cerezo, M., Arrasmith, A., Babbush, R., et al. (2021). Variational quantum algorithms. Nature Reviews Physics, 3, 625–644. 10.1038/s42254-021-00348-9

54. Abbas, A., Sutter, D., Zoufal, C., Lucchi, A., Figalli, A., & Woerner, S. (2020). The power of quantum neural networks. arXiv preprint arXiv:2011.00027. 10.48550/arXiv.2011.00027

55. Zeguendry, A., Jarir, Z., & Quafafou, M. (2023). Quantum Machine Learning: A Review and Case Studies. Entropy, 25(2), 287. 10.3390/e25020287

56. Gupta, S., & Deshmukh, N. D. (2022). Quantum computing to enhance performance of machine learning algorithms. In Apple Academic Press eBooks (pp. 165–178). 10.1201/9781003277217-13

57. Gill, S. S., Cetinkaya, O., Marrone, S., Claudino, D., Haunschild, D., Schlote, L., Wu, H., Ottaviani, C., Liu, X., Machupalli, S. P., Kaur, K., Arora, P., Liu, J., Farouk, A., Song, H. H., Uhlig, S., & Ramamohanarao, K. (2024, March 4). Quantum Computing: vision and challenges. arXiv.org. https://arxiv.org/abs/2403.02240

58. 58. Bowles, J., Ahmed, S., & Schuld, M. (2024, March 11). Better than classical? The subtle art of benchmarking quantum machine learning models. arXiv.org. https://arxiv.org/abs/2403.07059

59. Experimental benchmarking of quantum machine learning classifiers. (2023, November 8). IEEE Conference Publication | IEEE Xplore. https://ieeexplore.ieee.org/document/10343811

60. Chen, K.-C., Li, T.-Y., Wang, Y.-Y., See, S., Wang, C.-C., Wille, R., Chen, N.-Y., Yang, A.-C., & Lin, C.-Y. (2024). Validating large-scale quantum machine learning: Efficient simulation of quantum support vector machines using tensor networks. arXiv preprint arXiv:2405.02630. https://inspirehep.net/literature/2783664

61. Huang, H. Y., Broughton, M., Mohseni, M., et al. (2021). Power of data in quantum machine learning. Nature Communications, 12, 2631. 10.1038/s41467-021-22539-9

62. Liu, H. (2024). Research on Quantum Computing Acceleration of support vector machines in multi-dimensional nonlinear feature spaces. Applied and Computational Engineering, 100(1), 99–108. 10.54254/2755-2721/2025.17860

63. Gamble, S. (2019). Quantum computing: What it is, why we want it, and how we’re trying to get it. In National Academy of Engineering, Frontiers of Engineering: Reports on Leading-Edge Engineering from the 2018 Symposium. National Academies Press. https://www.ncbi.nlm.nih.gov/books/NBK538701/

64. Kavitha, S. S., & Kaulgud, N. (2024). Quantum machine learning for support vector machine classification. Evolutionary Intelligence, 17, 819–828. 10.1007/s12065-022-00756-5

65. Ahsan, M. M., Mahmud, M. A. P., Saha, P. K., Gupta, K. D., & Siddique, Z. (2021). Effect of Data Scaling Methods on Machine Learning Algorithms and Model Performance. Technologies, 9(3), 52. 10.3390/technologies9030052

66. Kumar, N., Tripathi, S., Sharma, N., Patiyal, S., Devi, N. L., & Raghava, G. P. (2024). A method for predicting linear and conformational B-cell epitopes in an antigen from its primary sequence. Computers in Biology and Medicine, 170, 108083. 10.1016/j.compbiomed.2024.108083

67. Vita, R., Mahajan, S., Overton, J. A., Dhanda, S. K., Martini, S., Cantrell, J. R., Wheeler, D. K., Sette, A., & Peters, B. (2018). The Immune Epitope Database (IEDB): 2018 update. Nucleic Acids Research. 10.1093/nar/gky1006

68. Bateman, A., Martin, M., Orchard, S., Magrane, M., Ahmad, S., Alpi, E., Bowler-Barnett, E. H., Britto, R., Bye-A-Jee, H., Cukura, A., Denny, P., Dogan, T., Ebenezer, T., Fan, J., Garmiri, P., Da Costa Gonzales, L. J., Hatton-Ellis, E., Hussein, A., Ignatchenko, A., Zhang, J. (2022). UniProt: the Universal Protein Knowledgebase in 2023. Nucleic Acids Research, 51(D1), D523–D531. 10.1093/nar/gkac1052

69. Arora, A., Patiyal, S., Sharma, N., Devi, N. L., Kaur, D., & Raghava, G. P. S. (2023). A random forest model for predicting exosomal proteins using evolutionary information and motifs. PROTEOMICS, 24(6). 10.1002/pmic.202300231

70. Li, W., & Godzik, A. (2006). Cd-hit: a fast program for clustering and comparing large sets of protein or nucleotide sequences. Bioinformatics, 22(13), 1658–1659. 10.1093/bioinformatics/btl158

71. Chaudhary, K., Kumar, R., Singh, S. et al. A Web Server and Mobile App for Computing Hemolytic Potency of Peptides. Sci Rep 6, 22843 (2016). 10.1038/srep22843

72. Gautam, A., Chaudhary, K., Singh, S., Joshi, A., Anand, P., Tuknait, A., Mathur, D., Varshney, G. C., & Raghava, G. P. S. (2013). Hemolytik: a database of experimentally determined hemolytic and non-hemolytic peptides. Nucleic Acids Research, 42(D1), D444–D449. 10.1093/nar/gkt1008

73. Bairoch, A., & Apweiler, R. (2000). The SWISS-PROT protein sequence database and its supplement TrEMBL in 2000. Nucleic Acids Research, 28(1), 45–48. 10.1093/nar/28.1.45

74. Rathore, A. S., Choudhury, S., Arora, A., Tijare, P., & Raghava, G. P. (2024). ToxinPred 3.0: An improved method for predicting the toxicity of peptides. Computers in Biology and Medicine, 179, 108926. 10.1016/j.compbiomed.2024.108926

75. Kaas, Q., Westermann, J.-C., Halai, R., Wang, C. K. L., & Craik, D. J. (2008). ConoServer, a database for conopeptide sequences and structures. Bioinformatics, 24(3), 445–446. 10.1093/bioinformatics/btm596

76. Shi, G., Kang, X., Dong, F., Liu, Y., Zhu, N., Hu, Y., Xu, H., Lao, X., & Zheng, H. (2021). DRAMP 3.0: an enhanced comprehensive data repository of antimicrobial peptides. Nucleic Acids Research, 50(D1), D488–D496. 10.1093/nar/gkab651

77. Waghu, F. H., Barai, R. S., Gurung, P., & Idicula-Thomas, S. (2015). CAMPR3: a database on sequences, structures and signatures of antimicrobial peptides: Table 1. Nucleic Acids Research, 44(D1), D1094–D1097. 10.1093/nar/gkv1051

78. Jhong, J., Yao, L., Pang, Y., Li, Z., Chung, C., Wang, R., Li, S., Li, W., Luo, M., Ma, R., Huang, Y., Zhu, X., Zhang, J., Feng, H., Cheng, Q., Wang, C., Xi, K., Wu, L., Chang, T., .. . Lee, T. (2021). dbAMP 2.0: updated resource for antimicrobial peptides with an enhanced scanning method for genomic and proteomic data. Nucleic Acids Research, 50(D1), D460–D470. 10.1093/nar/gkab1080

79. Piotto, S. P., Sessa, L., Concilio, S., & Iannelli, P. (2012). YADAMP: yet another database of antimicrobial peptides. International Journal of Antimicrobial Agents, 39(4), 346–351. 10.1016/j.ijantimicag.2011.12.003

80. Pirtskhalava, M., Amstrong, A. A., Grigolava, M., Chubinidze, M., Alimbarashvili, E., Vishnepolsky, B., Gabrielian, A., Rosenthal, A., Hurt, D. E., & Tartakovsky, M. (2020). DBAASP v3: database of antimicrobial/cytotoxic activity and structure of peptides as a resource for development of new therapeutics. Nucleic Acids Research, 49(D1), D288–D297. 10.1093/nar/gkaa991

81. Pande, A., Patiyal, S., Lathwal, A., Arora, C., Kaur, D., Dhall, A., Mishra, G., Kaur, H., Sharma, N., Jain, S., Usmani, S. S., Agrawal, P., Kumar, R., Kumar, V., & Raghava, G. P. (2022). PFeature: a tool for computing wide range of protein features and building prediction models. Journal of Computational Biology, 30(2), 204–222. 10.1089/cmb.2022.0241

82. Patro, S. G., & Sahu, K. K. (2015). Normalization: A preprocessing stage. *International Advanced Research Journal in Science*, Engineering and Technology (IARJSET), 2(3). 10.17148/IARJSET.2015.2305

83. Probst, P., Boulesteix, A.-L., & Bischl, B. (2019). Tunability: Importance of hyperparameters of machine learning algorithms. Journal of Machine Learning Research, 20(1), 1934–1965. https://dl.acm.org/doi/10.5555/3322706.3361994

84. Weerts, H. J., Mueller, A. C., & Vanschoren, J. (2020). Importance of Tuning Hyperparameters of Machine Learning Algorithms. *ArXiv*. https://arxiv.org/abs/2007.07588

85. Bergholm, V., Izaac, J., Schuld, M., Gogolin, C., Ahmed, S., Ajith, V., Alam, M. S., Alonso-Linaje, G., AkashNarayanan, B., Asadi, A., Arrazola, J. M., Azad, U., Banning, S., Blank, C., Bromley, T. R., Cordier, B. A., Ceroni, J., Delgado, A., Olivia, D. M., Killoran, N. (2018, November 12). PennyLane: Automatic differentiation of hybrid quantum-classical computations. arXiv.org. https://arxiv.org/abs/1811.04968

86. West, M. T., Sevior, M., & Usman, M. (2023). Boosted ensembles of Qubit and continuous variable quantum support vector machines for B Meson flavor tagging. Advanced Quantum Technologies, 6(10). 10.1002/qute.202300130

87. Almonteros, J. R., & Matias, J. B. (2024). Integration of stratified k-fold cross-validation to enhance prediction accuracy: A comparison study. 2024 *5th International Conference on Data Analytics for Business and Industry (ICDABI)*, 81–85. 10.1109/ICDABI63787.2024.10800425

88. 88. Berrar, D. (2018). Cross-Validation. In Elsevier eBooks (pp. 542–545). 10.1016/b978-0-12-809633-8.20349-x

89. Željko Ð. Vujovic, “Classification Model Evaluation Metrics” International Journal of Advanced Computer Science and Applications(IJACSA), 12(6), 2021. 10.14569/IJACSA.2021.0120670

90. Sharp, T. (2008). Implementing decision trees and forests on a GPU. In Lecture notes in computer science (pp. 595–608). 10.1007/978-3-540-88693-8_44

91. Ha, Y., Shashaani, S., & Menickelly, M. (2024). Two-Stage estimation and variance modeling for Latency-Constrained variational quantum algorithms. INFORMS Journal on Computing. 10.1287/ijoc.2024.0575

92. Abbas, A., Ambainis, A., Augustino, B. et al. Challenges and opportunities in quantum optimization. Nat Rev Phys 6, 718–735 (2024). 10.1038/s42254-024-00770-9

93. Zhuang, S., Tanner, J., Wu, Y., et al. (2024). Non-hemolytic peptide classification using a quantum support vector machine. Quantum Information Processing, 23, 379. 10.1007/s11128-024-04540-5

94. Havenstein, C., Thomas, D., & Chandrasekaran, S. (2018). Comparisons of performance between quantum and classical machine learning. SMU Data Science Review, 1(4), Article 11. Available at https://scholar.smu.edu/datasciencereview/vol1/iss4/11

95. Sciorilli, M., Borges, L., Patti, T.L. et al. Towards large-scale quantum optimization solvers with few qubits. Nat Commun 16, 476 (2025). 10.1038/s41467-024-55346-z

